# Characterization of novel TMEM173 mutation with additive IFIH1 risk allele

**DOI:** 10.1101/394353

**Authors:** Salla Keskitalo, Emma Haapaniemi, Elisabet Einarsdottir, Kristiina Rajamäki, Hannele Heikkilä, Mette Ilander, Minna Pöyhönen, Ekaterina Morgunova, Kati Hokynar, Sonja Lagström, Sirpa Kivirikko, Satu Mustjoki, Kari Eklund, Janna Saarela, Juha Kere, Mikko Seppänen, Annamari Ranki, Katariina Hannula-Jouppi, Markku Varjosalo

## Abstract

*TMEM173* encodes for STING that is a transmembrane protein activated by pathogen or self-derived cytosolic nucleic acids causing its translocation from ER to Golgi, and further to vesicles. Monogenic STING gain-of-function mutations cause early-onset type I interferonopathy, with disease presentation ranging from fatal vasculopathy to mild chilblain lupus. Molecular mechanisms causing the poor phenotype-genotype correlation are presently unclear. Here we report a novel gain-of-function G207E STING mutation causing a distinct phenotype with alopecia, photosensitivity, thyroid dysfunction, and STING-associated vasculopathy with onset in infancy (SAVI) -features; livedo reticularis, nasal septum perforation, facial erythema, bacterial infections and skin vasculitis. Single residue polymorphisms in *TMEM173* and an *IFIH1* T946 risk allele modify disease presentation in the affected multigeneration family, explaining the varying clinical phenotypes. The G207E mutation causes constitutive activation of inflammation-related pathways in HEK cells, as well as aberrant interferon signature and inflammasome activation in patient PBMCs. Protein-protein interactions further propose impaired cellular trafficking of G207E mutant STING. These findings reveal the molecular landscape of STING and highlight the complex additive effects on the phenotype.

**BRIEF SUMMARY:** Novel gain-of-function mutation in *TMEM173,* associated with single residue polymorphisms in *TMEM173* and *IFIH1*, causes a distinct clinical phenotype with some shared features of SAVI.

## Introduction

Monogenic interferonopathies are characterized by increased type I interferon signaling leading to vasculopathy, autoinflammation and systemic lupus erythematosus (SLE)-like disease (1). In these, increased sensing of endo- and exogenous nucleic acids leads to hyperactive Toll-likereceptors (TLRs), RNA and DNA sensors, and to an enhanced type I interferon (IFN) response (2). The upregulation of type I IFNs is thought to maintain the persistent self-directed immune reaction (3).

One of the 18 currently known monogenic interferonopathies is STING-associated vasculopathy with onset in infancy (SAVI; MIM 615934) that is caused by mutations in *TMEM173* (4, 5). *TMEM173* encodes for the stimulator of interferon genes (STING), a transmembrane protein residing in the endoplasmic reticulum (ER). It has an essential role in sensing cytosolic double stranded DNA (dsDNA) and directly binding to bacterial second messengers, such as cyclic dinucleotides (CDNs) c-di-GMP, c-di-AMP and 3’3’-cGAMP (6, 7). The recognition of nucleic acids or cyclic nucleotides initiates the production of type I IFN and other inflammatory cytokines leading to nucleicacid driven inflammation.

STING has four amino-terminal transmembrane domains spanning the first 136 amino acid residues, followed by the helix α1 at residues 153-177 (8). Helix α1, also known as the dimerization domain, is essential for protein stability, intraprotein interactions and ligand binding (9). The CDN binding domain (residues 153-340) is part of the cytoplasmic carboxy-terminus, stretching over residues 138-379 and having multiple phosphorylation and downstream signaling interaction sites (8, 10).

The first described constitutively active *TMEM173* mutations are situated at or near the hydrophobic helix α1 dimerization domain and associated with early-onset vasculitis, autoinflammation and interstitial lung disease that define the SAVI phenotype (5, 11, 12). A recent study identified five patients with a GOF mutation affecting the dimerization domain (13). In contrast to the severely affected infants (5) these patients only presented with mild skin vasculitis and were diagnosed with familial chilblain lupus (13). In addition to the mutations in the dimerization domain, proximal substitutions affecting the CDN binding domain were reported in single patients presenting with variable phenotypes of STING-associated autoinflammation (14, 15). Overall, all of the reported *TMEM173* mutations have been GOF, leading to elevated IFNB production which activates the JAK/STAT-pathway and generates a positive feedback loop (16). As there is poor correlation between genotype and clinical phenotype, additional, poorly understood intrinsic or environmental factors likely modify the disease outcome. Another important interferonopathy gene is the IFN-induced helicase C domain-containing protein 1 (*IFIH1,* also known as MDA5). IFIH1 functions as a nucleic acid sensor in the cytoplasm, inducing transcription of type I IFN and IFN-regulated genes upon activation. GOF *IFIH1* variants are seen in Aicardi-Goutières and Singleton-Merten syndromes, both of which feature prominent vascular inflammation (17-19). Also, polymorphism in *IFIH1* has been linked to SLE (rs1990760, p.A946T) (20, 21) leading to a variable phenotype spectrum (22). The A946T GOF risk variant leads to increased production of type I IFN, promoting inflammation and elevating the risk of autoimmunity. The T946 allele also modifies the effects of other autoimmune risk alleles, which leads to variable disease severity (23, 24).

Here we report a multigenerational family of six affected individuals, presenting with several lupoid-like features (photosensitivity, alopecia, autoimmune thyroiditis) and features of SAVI (bacterial skin infections, early-onset livedo reticularis, nasal septum perforation and facial erythema). Affected individuals have a combined genotype of (1) a novel disease-causing *TMEM173* mutation G207E together with (2) either arginine or histidine at a polymorphic position 232 in *TMEM173*, and (3) homo/heterozygosity for the *IFIH1* T946 risk allele. We describe the additive effects of these multiplexed single residue polymorphisms using cell models and patient material, highlighting the effect of common genetic variation in rare disease presentations. The results broaden the spectrum of published *TMEM173* mutation-associated phenotypes, provide insight into the activation of alternative NLRP3 inflammasome and reveal the STING interactome.

## Results

### Clinical presentation and immune phenotype

We evaluated a multigenerational Finnish family with a few SAVI-like features, but presenting also multiple symptoms previously not associated with SAVI or chilblain lupus (**Supplementary figure 1A)**. Of 10 affected family members, six were available for detailed clinical evaluation (**Figure 1A**) and the main clinical findings are summarized in **Table 1**. Affected family members presented with novel STING mutation induced features of photosensitivity, alopecia ranging from areata to universalis and autoimmune thyroiditis. Prominent features included early-onset persistent livedo reticularis, skin vasculitis, recurrent and severe infections variably reported in SAVI (**Figure 1B-E**). None of the patients had typical SAVI associated interstitial lung disease, violaceus facial, nasal, auricular or acral patches progressing to ulcers or necrosis, febrile attacks, failure to thrive, elevated inflammatory markers or autoantibodies **(Supplementary figure 1B** and **Supplementary table 1**). The most severely affected family members suffered from UV exposure induced cutaneous vasculitis, nasal septum perforation, severe infections including necrotizing cellulitis and abscesses concomitant with SAVI (**Table 1** and **Supplementary figure 1B).**

**Table 1:** Genotypes and clinical features of affected family members.

| Patient demographics |  |  |  |  |  |  |
| --- | --- | --- | --- | --- | --- | --- |
| Patient | IV.1 | V.2 | III.4 | III.9 | III.7 | IV.6 |
| Age (years) | 33 | 12 | 52 | 49 | 56 | 31 |
| Gender | male | female | female | female | female | female |
| STING G207E | G/E | G/E | G/E | G/E | G/E | G/E |
| STING R232H rs1131769 | H/R | H/R | H/R | H/H | H/H | H/R |
| IFIH1-risk allele A946T rs1990760 | A/T | T/T | A/T | A/T | A/A | A/A |
| Inflammatory manifestations |  |  |  |  |  |  |
| Livedo reticularis | Widespread birth | Widespread birth | Localized birth | Widespread adulthood | Widespread 8y | Localized early childhood |
| Skin vasculitis | yes 29 y | yes 10 y | no | no | no | no |
| Nasal septal perforation | yes 29 y | yes 6 y | no | yes 49 y | no | no |
| Facial erythema | yes | yes | yes | yes | yes | no |
| UV sensitivity | burns easily | burns easily | burns easily | urticaria | burns easily | burns easily |
| Alopecia at age of years | universalis 10 y | universalis 6 y | areata 7 y | totalis 4 y (grew back) and universalis 30 y | no | universalis 10 y (scalp hair grew back) and universalis 12 y |
| Thyroid dysfunction | no | hypo 12 y | hypo 7y | hyper 38 y | no | hyper 13 y |
| Autoimmune thyroiditis | no | yes | no | yes | no | yes |
| Infection susceptibility |  |  |  |  |  |  |
| Skin infections | facial erysipelas 33 y, deep abscess in right thigh 27y | necrotizing fascitis and abscesses in upper extremity 3 y, necrotic vasculitis and abscess in the abdomen 10 y | no | no | abscess in left inguinum childhood | no |
| Periodontitis | no | no | no | no | yes 42 y | yes 30 y |
| Respiratory tract | no | pneumonia 1.5 y | no | recurrent sinusitis adulthood | recurrent sinusitis adulthood | no |

**Figure 1:**
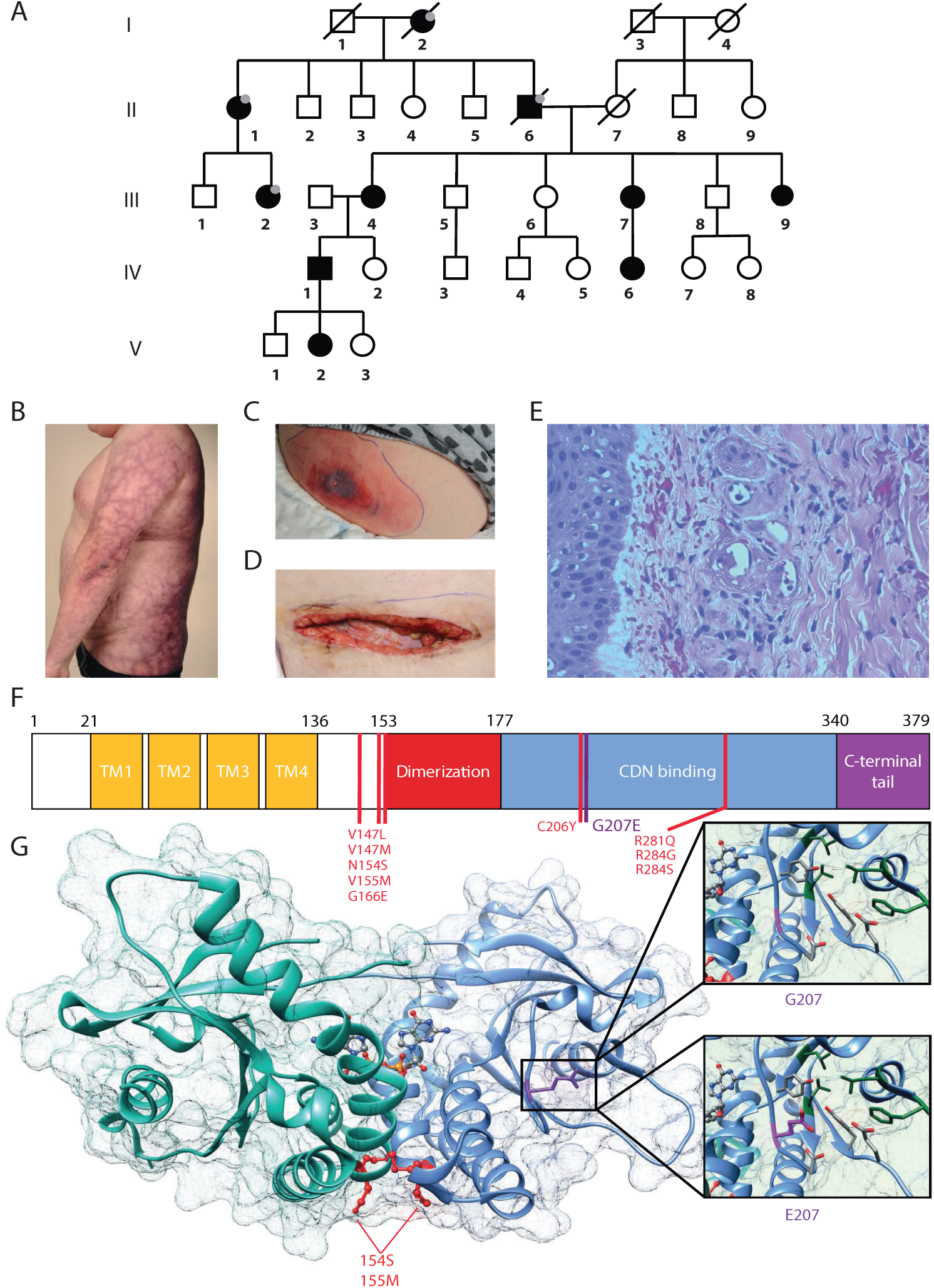
G207E STING mutation associates with SAVI and lupoid-like features. Family pedigree. Individuals without in-depth clinical evaluation are denoted with a gray dot; deceased individuals by diagonal bars. B) Livedo reticularis in IV.1. C-D) Necrotizing cellulitis and vasculitis in V.2 at initial situation (C) and after surgical revision (D). E) Skin biopsy from IV.1 reveals vasculitis with thrombotic and destructed blood vessels with neutrophils in their walls as well as thickened capillaries surrounded by inflammatory cells and extravasated erythrocytes. F) Schematic representation of the STING protein showing transmembrane (TM), dimerization, and cyclic-di-nucleotide (CDN) binding domains together with the carboxy-terminal tail. Previously identified human gain-of-function mutations are denoted in red and the here identified G207E mutation is shown in purple. G) Crystallographic structure of STING dimer in complex with cGAMP, featuring dimerization and c-di-GMP binding domains (PDB entry 4EMT). Glutamine (upper panel) contains no side chain and cannot contact cGAMP or polar (red) or hydrophobic (green) residues nearby. Glutamic acid, in contrast, has a polar, highly flexible side chain that can rotate and interact with polar amino acids and cGAMP (left). SAVI mutations are shown in red at the bottom of the dimer.

All six evaluated patients had increased peripheral CD4+ T cell and naïve CD4+CCR7^+^CD45RA^+^ T cell count (Table 2). The central memory CD4+ T cells numbers remained unaffected, but pronounced decreases were observed in the effector memory subset. These observations are in accordance with recent findings on patients carrying activating *TMEM173* mutations (25). Among B cell populations, both naïve (CD27^-^IgD^+^) and immune-exhausted activated (CD38^low^CD21^low^) B cells were increased in 4/6 patients (Table 2). Immunologic workup of these six patients further showed normal acute phase reactants, blood cell counts, various autoantibodies, complement activation (Supplementary table 1), proliferative lymphocyte responses to mitogens and antibody responses to both protein and polysaccharide vaccines (data not shown).

**Table 2:**
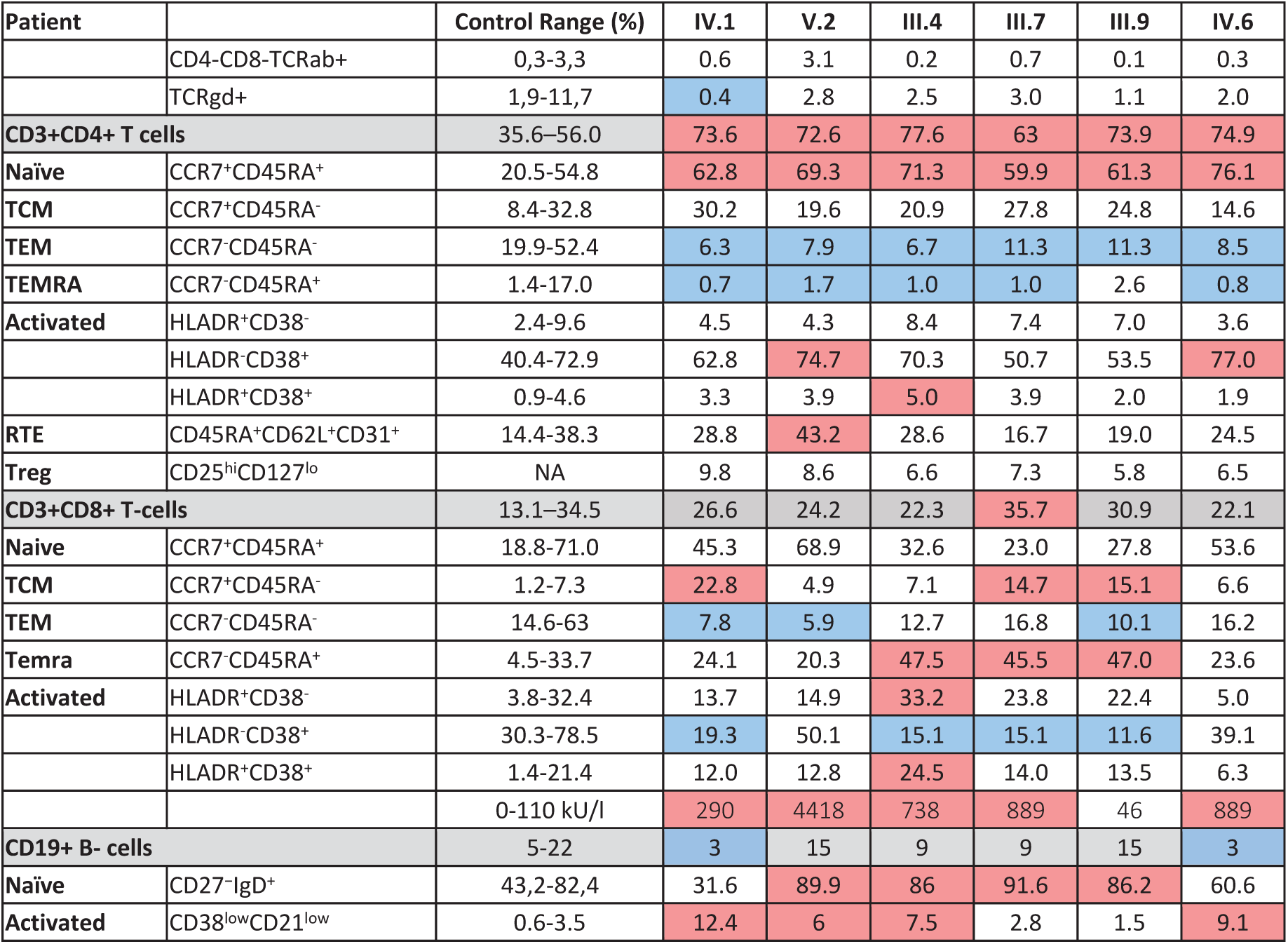
Hematologic parameters of Tand B-cells of affected family members. Values higher than control range are highlighted with red, whereas lower values are indicated with blue background.

### Identification and structural analysis of TMEM173 mutation

Based on the pedigree (**Figure 1A**), we assumed an autosomal dominant inheritance. We performed linkage analysis on 12 unaffected and 6 affected family members and whole genome sequencing on patients IV.1 and V.2. Considering all affected, Merlin non-parametric linkage analysis revealed no statistically significant linkage peaks. NPL scores of >1.7 were observed at four loci on chromosomes 3, 5, 9, and 12. In these, only one co-segregating rare variant negatively affected an evolutionary conserved residue. This heterozygous missense mutation Chr5 (GRCh38): g.139478409C>T results in a p.Gly207Glu (G207E) substitution in *TMEM173* encoding for STING and was absent from major public and in-house genome databases. It localizes next to a previously described mutation p.Cys206Tyr (C206Y) (14) lying within the CDN binding domain. The G207E mutation locates in a fold that participates in substrate binding and STING folding, and changes the electrically neutral glycine to negatively charged glutamic acid (**Figure 1F-G**). Based on the protein structure prediction, the G207E mutation is expected to change the electric charge of the protein fold, destabilizing it and thus potentially altering its function (**Figure 1G** and **Table 3**). For further structural comparison of C206Y and G207E, we predicted the effects of single residue changes on protein structure and stability using Site Directed Mutator (SDM) (26). SDM predicted that both C206Y and G207E reduce the stability of the protein (pseudo ΔΔG -1.27/-1.28 and-0.72/-0.63, respectively **Table 3**) and likely lead to protein malfunction and disease. The C206Y mutation with newly acquired aromatic side chain results in more drastic structural alteration than G207E. Interestingly, polymorphism at position 232 in *TMEM173* influenced G207E’s relative side chain solvent accessibility suggesting greater ligand accessibility and binding with variant R232 compared to H232. C206Y, as a buried residue, remained solvent inaccessible independent of position 232.

**Table 3:** Predictions on protein stability with C206Y and G207E site mutagenesis. Percentage relative sidechain solvent accessibility (RSA) for wildtype (WT-RSA %) and both mutant (MUT-RSA %) residues at 206 and 207. Residue depth for wildtype (WT-DEPTH) and mutants (MUT-DEPTH) is presented in angstrom units (Å). Residue occluded surface packing is likewise presented for wildtype (WT-OSP) and mutant (MUT-OSP) residues. Predicted pseudo ΔΔG and its effect on protein stability (outcome) are presented.

| Mutation | PDB File | Residue 232 | WT-RSA (%) | WT-DEPTH (Å) | WT-OSP | MUT-RSA (%) | MUT-DEPTH (Å) | MUT-OSP | Predicted pseudo $\Delta\Delta G$ | Outcome |
| --- | --- | --- | --- | --- | --- | --- | --- | --- | --- | --- |
| C206Y | 4LOI | H | 0.0 | 7.4 | 0.56 | 0.0 | 8.0 | 0.7 | -1,27 | Reduced stability |
| G207E | 4LOI | H | 57.1 | 4.4 | 0.35 | 49.8 | 3.5 | 0.27 | -0,72 | Reduced stability |
| C206Y | 4KSY | R | 0.0 | 6.7 | 0.53 | 0.1 | 7.0 | 0.64 | -1,28 | Reduced stability |
| G207E | 4KSY | R | 82.5 | 3.9 | 0.32 | 73.6 | 3.3 | 0.23 | -0,63 | Reduced stability |

### Identification of disease-modifying polymorphisms in TMEM173 and IFIH1

A non-synonymous variant of human STING encoding arginine (R) at position 232 has been identified as a major allele in the general population, in contrast to the histidine (H) 232 commonly listed as the wildtype allele (6). In accordance with the population data, the majority of the affected family members were heterozygotes for major allele R232 (**Table 1**). Additionally, single nucleotide polymorphism (rs1990760, Ala946Thr (A946T)) in *IFIH1* causes constitutively active IFN signaling and predisposition to autoimmune diseases (20, 23, 24). In the affected family 4/6 analyzed members had at least one copy of the *IFIH1* T946 risk allele (**Table 1**). When the clinical presentations of affected family members were correlated with their combined genotypes of the monogenic *TMEM173* mutation G207E, and R232H variants, as well as the presence of *IFIH1* T946, a trend of additive effect jointly promoting disease was observed from 207E/H232/AA towards 207E/ R232/TT (**Table 1**).

### G207E mutation in STING constitutively activates IFN-β, STAT1/2, STAT3 and NFKB transcription

To study the effect of STING mutations on activation of IFNB, STAT1/2, STAT3 and NFKB pathways, we utilized luciferase-based reporter assays specific for each pathway. The constitutively active STING p.Ans154Ser (N154S) SAVI mutant was used as a positive control in all the assays (5). To mimic the combined genotypes of the affected family members, we combined the G207E mutation either with variant R232 (major population allele and genotype of patients IV.1, V.2, III.4, IV.6; herein R+207E) or H232 (minor allele and genotype of patients III.9, III.7; herein H+207E). Both combinations clearly induced IFNB, STAT1/2 and STAT3 pathway activation, even in basal unstimulated state (**Figure 2A-C**), thus indicating 207E as a clear GOF mutation.

**Figure 2:**
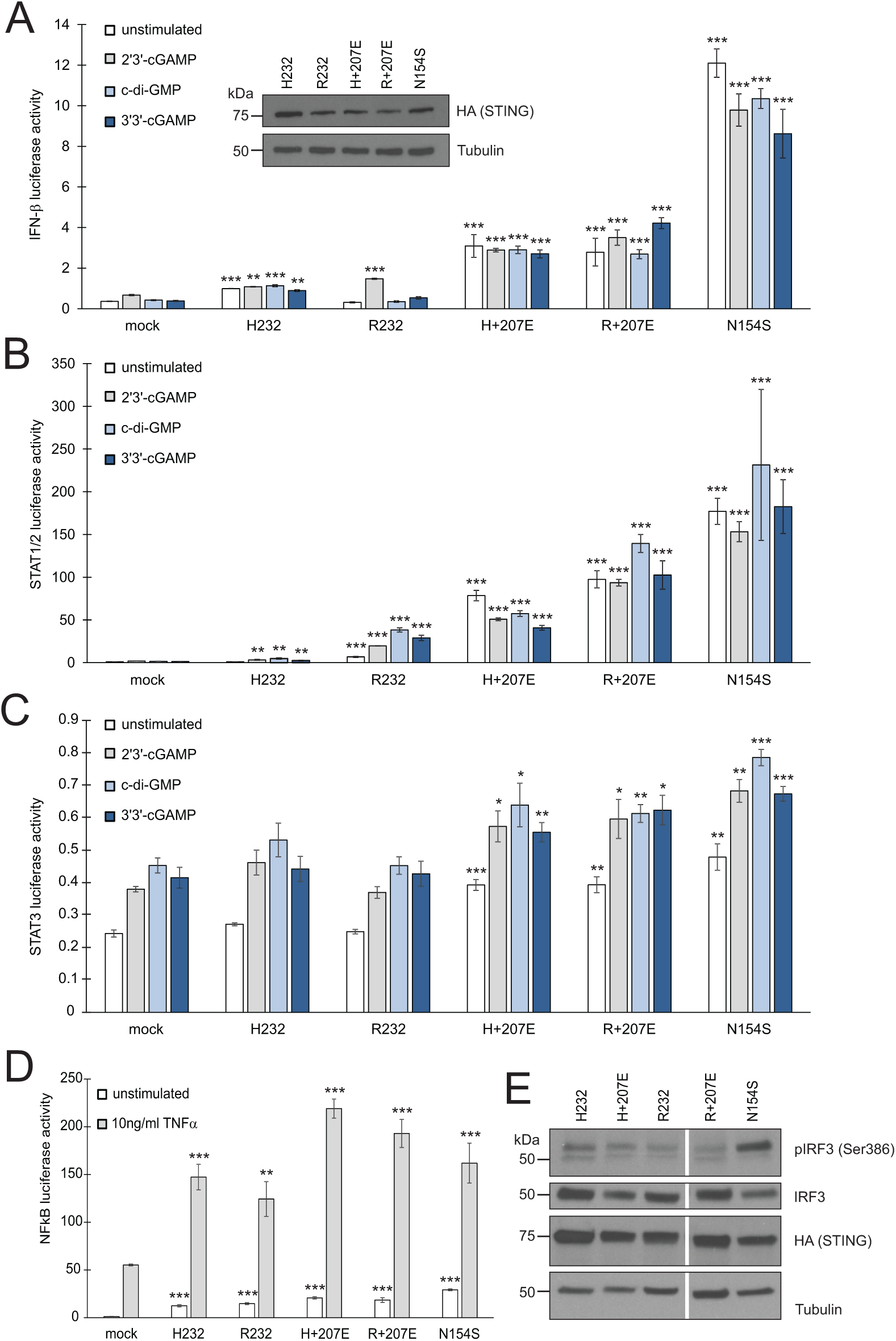
G207E constitutively activates several signal transduction pathways in HEK293 cells. The ratio of firefly luciferase to Renilla luciferase was calculated to analyze the fold induction of A) *IFNB* reporter, B) *STAT1/2*, C) *STAT3* or D) *NFKB* reporter after transient transfection either with empty vector (mock), H232, R232, H+207E, R+207E or SAVI mutant N154S. Expression of all the construct was verified by anti-HA antibody. Through A-C, the cells were stimulated with CDNs at concentration 4 µg/ml for 24 hours or left unstimulated. For *NFKB* reporter assay, the cells were stimulated with 10 ng/ml TNFA for 16 hours. In all the figures A to D mean of minimal three replicates together with SEM is presented. Statistical significance is indicated with asterisk (* p<0.05, ** p<0.01, *** p<0.001; paired t-test). E) The baseline levels of IRF3 phosphorylation remained unaltered in H232, H+207E, R232, R+207E. Only N154S presented elevated phosphorylation of IRF3.

The patients were susceptible to bacterial infections, but only a few remembered ever having viral infections such as common cold. Thus, we stimulated the transfected HEK293 cells with 2’3’cGAMP mimicking the mammalian response to cytosolic viral DNA, in addition to bacterial second messengers c-di-GMP and 3’3’-cGAMP (6, 27). After CDN stimulations variant R+207E presented slight increases of pathway activity in IFNB and STAT1/2 assays, while variant H+207E remained at constant level (IFNB) or showed diminished activity (STAT1/2) (**Figure 2A-B**). Both variants induced similar STAT3 and NFKB pathway activation upon stimulation (**Figure 2C-D**). As constitutive activation of STING has been thought to lead to IRF3 phosphorylation and subsequent expression of IFN-response genes (5, 28-30), we next analyzed IRF3 (Ser386) phosphorylation in stably expressing HEK293 cells (**Figure 2E**). Interestingly, IRF3 phosphorylation remained at similarly low levels in all studied combinations and their single counterparts, while N154S showed markedly elevated phosphorylation (**Figure 2E**). In concordance with our data, IRF3 independent disease development was recently reported in a STING N153S mouse model (31), demonstrating the involvement and activation of multiple pathways in STING-associated disease.

These results further confirm the crucial role of R at position 232 in cGAMP binding and IFN induction (32, 33), and show the constitutive activation of STING signaling pathways together with ligand-dependent hyperactivation of NFKB-pathway in G207E mutant cells.

### Baseline expression of interferon signature genes and JAK/STAT pathway components are elevated in patients with G207E mutation

To directly analyze baseline gene expression in patient PBMCs, we designed a custom Nanostring (34) gene set targeting multiple IFN-inducible and IFN signaling-related genes, as well as genes encoding components of the NFKB-regulated inflammasome pathway (**Figure 3A**). The IFN pathway-related target genes were selected based on their involvement in SLE, SAVI and familial chilblain lupus (5, 13, 35, 36). Strong upregulation of IFN-regulated genes was detected in an unrelated Singleton-Merten-patient with dominant GOF *IFIH1* mutation, validating the gene set and analysis pipeline (**Figure 3A**). The severely affected G207E patients carrying the *IFIH1* risk allele (IV.1, V.2, III.4) displayed higher IFN-regulated mRNA levels (marked in red) than the patient with G207E mutation alone (III.7) (**Figure 3A**).

**Figure 3:**
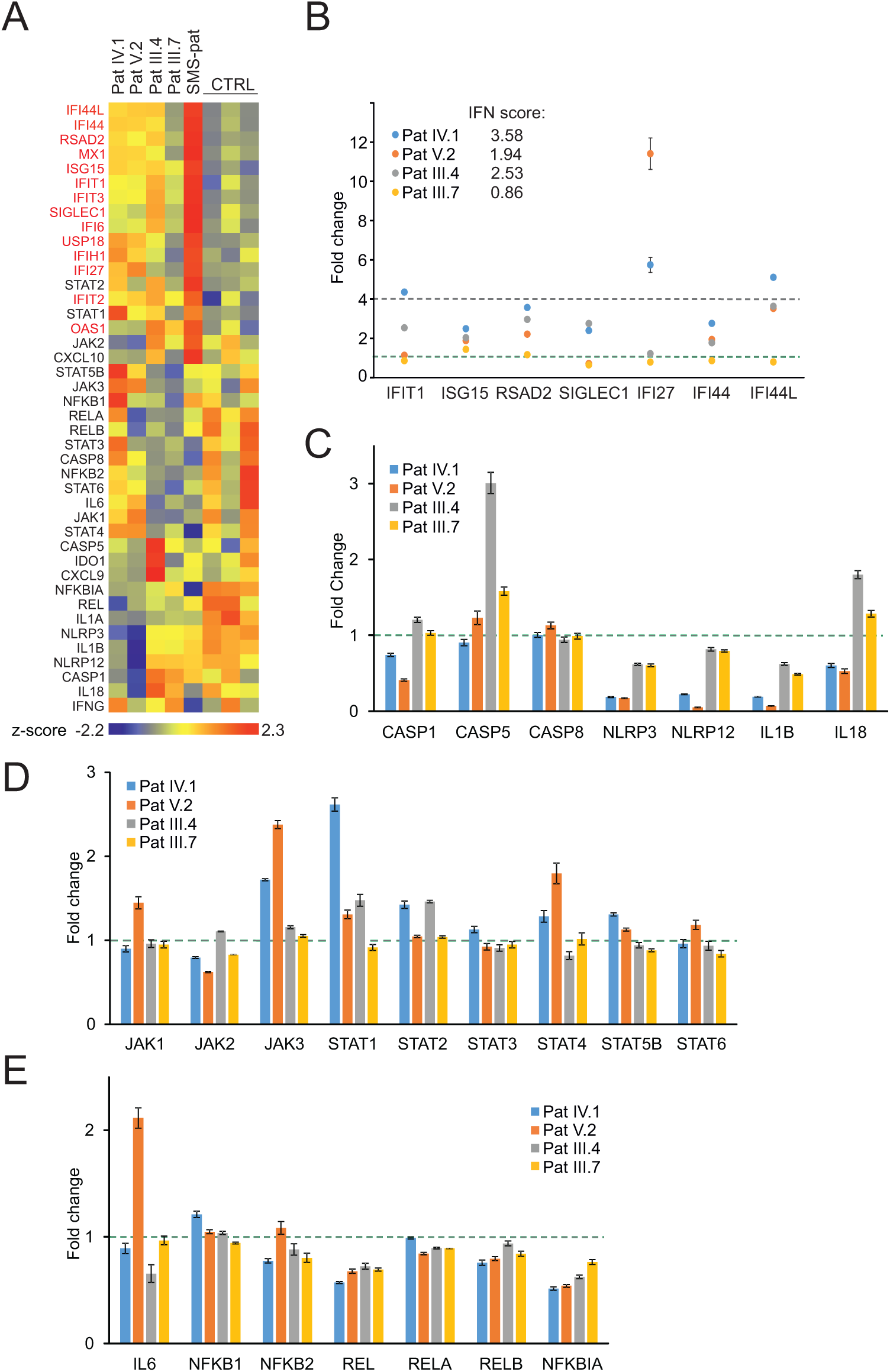
Gene expression analysis of patient PBMCs reveals alterations in IFN-regulated genes and multiple signaling pathways. A) Heat map of baseline gene expression of a custom Nanostring gene set in PBMCs of four G207E patients, a Singleton-Merten syndrome patient (SMS-pat; another type I interferon-mediated disease), and three healthy controls. After background thresholding and normalization, agglomerative clustering was performed on z-scores of genes. IFN-regulated genes are marked in red. B) mRNA fold changes of seven IFNregulated genes (*IFIT1*, *ISG15*, *RSAD2*, *SICLEC1*, *IFI27*, *IFI44*, and *IFI44L*) and the resulting IFN score for each patient. The values were normalized to internal controls and housekeeping genes. C-E) Fold changes in unstimulated patient PBMCs of inflammasome (C), JAK/STAT pathway (D), and NFkB pathway (E) related genes. Fold change 1 and 4 are indicated with green and blue dotted lines, respectively. Through A to E mean of three measurements together with SD is presented.

For IFN signature analysis (**Figure 3B**) we selected seven genes (*IFIT1*, *ISG15*, *RSAD2*, *SIGLEC1*, *IFI27*, *IFI44*, and *IFI44L*) based on previous publications on IFN signature in type I interferonopathies (13, 36). For fold change calculation, the patient data was normalized to matched control. We noted pronouncedly different gene expression pattern and elevated IFN scores in patients with G207E mutation and *IFIH1* risk allele (**Figure 3B**). The patient without IFIH1 A946T variant did not show such a drastic response, but scored above the healthy control threshold (36).

The patient cells displayed no clear overall tendency of baseline mRNA levels of inflammasome-related genes, whereas JAK/STAT-pathway components that mediate IFN signaling were upregulated, and selected NFKB-related genes, including the key negative regulator NFKBIA, were downregulated (**Figure 3C-E**). The trend of additive effects of genetic variance was most pronounced with STING downstream targets JAK1, JAK3, STAT4, STAT6, and IL6, where patient V.2 presented the highest baseline gene expression levels (**Figure 3D-E**).

Enhanced type I interferon response and aberrant inflammasome activation in patients with G207E mutation

STING has been recently linked with activation of the NLRP3 inflammasome (37, 38) that controls the proteolytic maturation and secretion of cytokines IL1B and IL18. In monocytes the NLRP3 inflammasome can be activated either by the canonical, non-canonical or alternative pathway (39). To test whether inflammasome was hyperactive in the patients, we stimulated their PBMCs with LPS and a subsequent ATP pulse to activate the canonical inflammasome or with prolonged LPS alone to trigger the alternative pathway. The secretion of IL1B and IL18 was used as a proxy for inflammasome activation. Pam3Cys that activates NFKB pathway via Toll-like receptor (TLR) 1/2, but does not normally trigger inflammasome signaling or type I IFN response, was used as a control (40). In the patients’ PBMCs LPS induced secretion of mature IL1B and IL18 via the alternative NLRP3 inflammasome pathway (**Figure 4A-B**) and, importantly, IL1B secretion was aberrantly triggered by Pam3Cys (**Figure 4A**).

**Figure 4:**
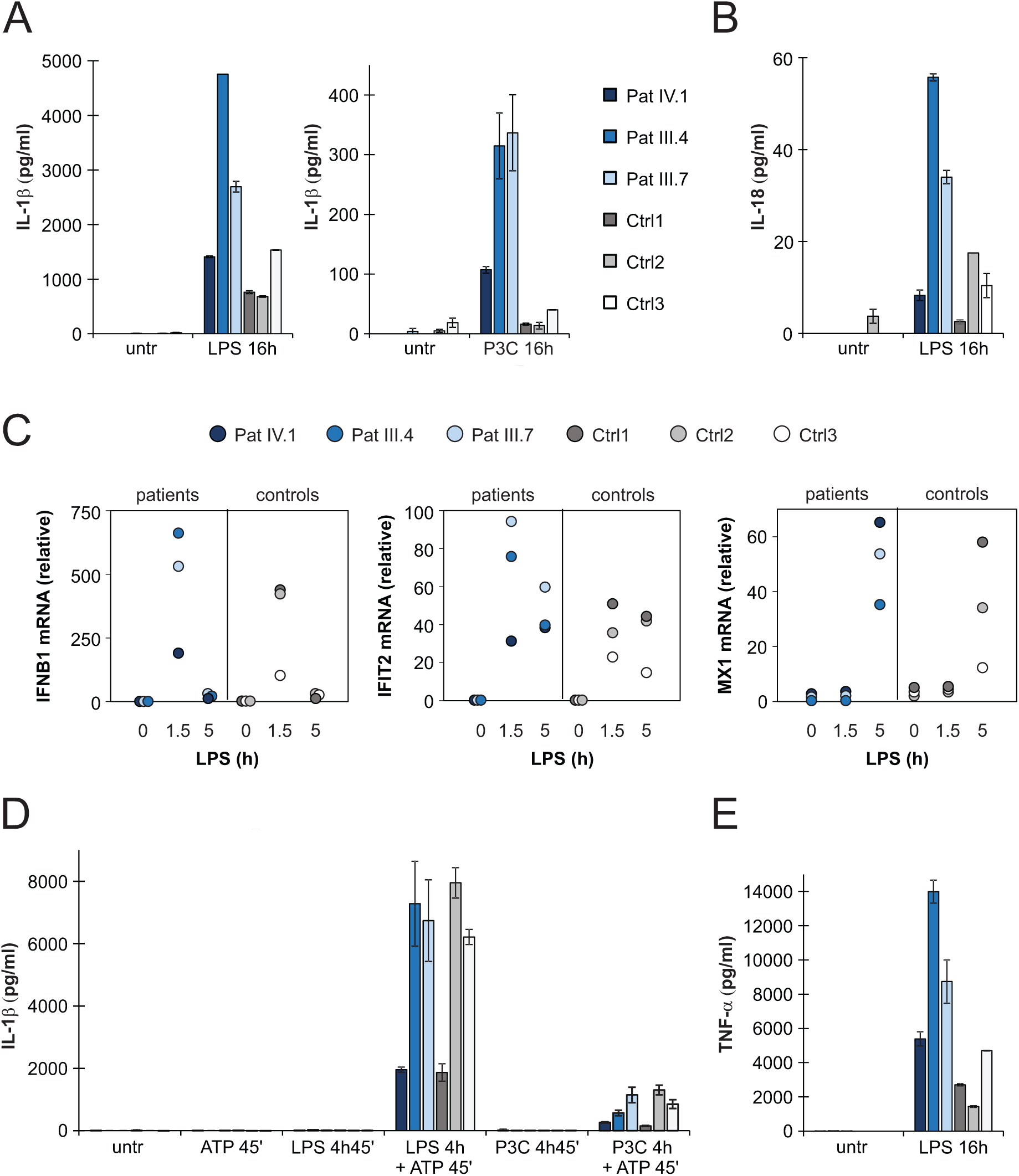
Type I interferon response and NLRP3 inflammasome activation in patient PBMCs. A-B) Secretion of IL1B and IL18 in PBMC culture supernatants after activation of the NLRP3 inflammasome via the alternative pathway using prolonged LPS or Pam3Cys (P3C) stimulation. C) RT-PCR analysis of *IFNB1*, *IFIT2* and *MX1* mRNAlevels in patient PBMCs left untreated and after LPS treatment. D) Secretion of IL1B in PBMC supernatant after canonical inflammasome pathway activation with LPS or Pam3Cys priming followed by ATP stimulation. E) TNFA concentration in PBMC culture supernatants after LPS stimulation. The data collection in A, B, D, and E was performed with enzyme-linked immunosorbent assay (ELISA). Data are presented from three patients and three control subjects. Cytokine levels represent means and the error bars denote SD from 2 biological replicates per condition; for RNA isolation and qPCR, the cells from these replicates were pooled.

The activation of alternative NLRP3 inflammasome in monocytes involves TLR4-TRIF signaling that also mediates LPS-induced type I IFN response (40). Thus, we analyzed the expression of IFNB and selected IFN-response genes after LPS treatment in patients’ PBMCs by RT-PCR that revealed elevated mRNA levels compared to controls (**Figure 4C)**. We also detected increases in pro-IL1B mRNA after LPS and Pam3Cys stimulation that may contribute to the elevated IL1B secretion, while mRNA levels of NLRP3 inflammasome components and IL18 did not show consistent alterations (**Supplementary figure 2A-B**). However, the patients’ PBMCs could not release mature IL1B in response to ATP alone without pre-priming with LPS or Pam3Cys to induce pro-IL1B transcription (**Figure 4D**). These results indicate that the patient genotype results in aberrant TLR-induced inflammasome responses mainly via increased NLRP3-mediated proteolytic maturation of the inflammasome target cytokines, rather than via increased cytokine mRNA expression.

Since the patient responses to ATP stimulation after short priming were normal (**Figure 4D and Supplementary figure 2C**), we conclude that the mutant STING mainly heightens the alternative NLRP3 inflammasome signaling. As the patients also showed elevated TNFA response to LPS stimulation, the dysregulation of TLR responses likely extends beyond the alternative inflammasome pathway (**Figure 4E and Supplementary figure 2B**). However, we did not see a general correlation between clinical disease severity and cytokine expression (**Figure 4A-B, E**).

### Molecular context of mutant STING

To further characterize the molecular context of STING at near physiological levels (41), we analyzed the interaction profiles of H232 (wild-type; wt), I200N (null (42, 43), N154S (5), and H+207E in inducibly expressing HEK293 cell lines using BioID proximity labeling. The BioID-method tags the protein of interest (bait) with a modified biotin ligase, which adds biotin to the proximal proteins (preys) (44). The biotinylated and thus closely interacting proteins are further affinity-purified and analyzed with quantitative mass spectrometry. In line with previous data (33), the wt STING preferentially interacted with Golgi, ER, endosomal and mitochondrial proteins (**Figure 5A**). All studied STING mutants (I200N, G207E and N154S) had lost multiple protein-protein interactions compared to the wt STING. By using hierarchical clustering, the interactions of the mutants form a continuation from the null allele (I200N) to GOF SAVI mutant (N154S). The G207E mutation clusters between the wt and the SAVI mutant (**Figure 5A**). This result is consistent with the reporter assay data and milder clinical phenotype in G207E carriers (**Figure 2**).

**Figure 5:**
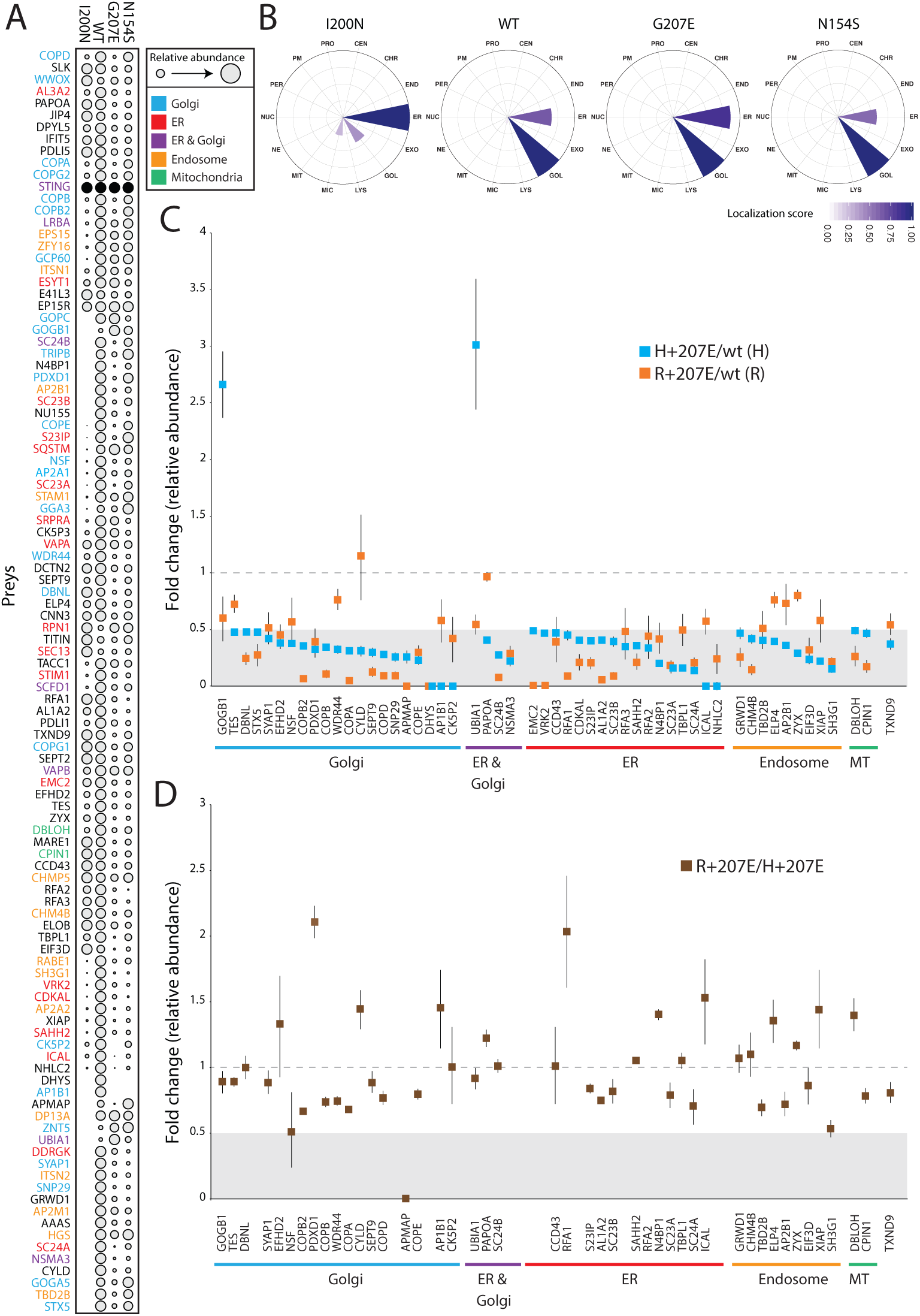
Functional data of STING wt and mutant cell lines shows differences in genotypes. A) Identified interacting proteins (preys) and their relative amount in stable I200N, H232 (wt), G207E, and N154S MAC-tagged Flp-In T-REx 293 cells. The relative amount of each prey was calculated after normalization to the bait expression. Dot size is indicative of relative prey abundance. B) Mass spectrometry microscopy assigned STING mutant and wt localizations at steady state. The circle indicates polar plots of 14 equally divided sectors of localization information. Localization abbreviations denote the following cellular compartments: peroxisome (PER), microtubule (MIC), endosome (ED), proteasome (PRO), nuclear envelope (NE), Golgi (GOL), lysosome (LYS), nucleolus (NUC), plasma membrane (PM), endoplasmic reticulum (EM), mitochondria (MIT), centrosome (CEN), chromatin (CHR), and exosome (EXO). Localization score is indicated with the color gradient. C) Comparison of identified proteins and their fold changes in H+207E and R+207E and their respective wild-type counterparts. D) Comparison of fold changes of R+207E to H+207E reveal genotype specific protein-protein interactions. In C and D light grey areas in figures indicate fold change 0-0.5, and dotted line fold change equal to 1.

These observations led us to utilize mass spectrometry microscopy (MS microscopy) (45) on our steady state BioID data of wt and mutant STING. The resolution of MS microscopy outperforms fluorescent microscopy in refining the exact subcellular molecular location (45). MS microscopy showed I200N localization mainly to ER, while G207E, N154S and wt were predominantly localized to Golgi (**Figure 5B**). Fascinatingly, the G207E localized more to the ER than wt or N154S (localization scores 0.86, 0.66 and 0.58, respectively) (**Figure 5B**). As localization score reflects the subcellular localization and the time spent there, we speculate that the constantly highly active N154S mutant rapidly translocates from the ER to Golgi (lowest ER localization score) leading to hyperactivation of downstream signaling. Meanwhile, the G207E is less active, translocates more slowly, and spends more time in the ER.

To investigate the G207E mutation more closely, we analyzed its interactomes in both allelic backgrounds (H+207E and R+207E). Generally, the G207E mutation led to a reduction of proteinprotein interactions with the proteins residing in Golgi, ER, endosome and mitochondria, compared to the corresponding wt (**Figure 5C**). Notably, the G207E mutant had weak interactions with all components of the coatomer complex I (COPI)machinery mediating retrograde transport from Golgi to the ER, and most strikingly with coatomer subunits A, B, B2, D, and E (COPA, COPB, COPB2, COPD, and COPE, respectively) (46, 47) (**Figure 5A, C**). Also, multiple components of the coatomer complex II responsible for ER to Golgi transport, such as syntaxin 5 (STX5) and protein transport protein Secs were downregulated (Sec23A, Sec23B, Sec24A, Sec24B) (**Figure 5A, C**). The interactions between GOGB1 and UBIA1 were stronger in the H+207E mutant (> 2-fold). UBIA1 localizes to ER-Golgi compartment, while GOGB1 mainly resides at the cis-Golgi participating in COPI-mediated transport (48-50). These data suggest that in addition to amplifying downstream signaling, the GOF mutations cause broad alterations in STING subcellular localization and function in the vesicular system, and reflect a plausible trafficking defect. The features overlap with other known monogenic autoimmune diseases that are caused by defects in the vesicular trafficking (51, 52). The alterations likely contribute to the autoimmune and infectious manifestations seen in the affected patients.

The clinical presentations suggested that the allelic variance at position 232 modifies the disease severity; therefore, we compared the protein interactions between variants R+207E and H+207E. These showed highly similar interac-tomes, with ∼2-fold differences in five interactions (**Figure 5D**). These genotype-specific interactions included lost association between R+207E and APMAP (adipocyte plasma membrane-associated protein with uncharacterized function), and increased interaction with PDXD1 (pyridoxal-dependent decarboxylase domain-containing protein 1 residing at Golgi) and RFA1 (Replication protein A 70 kDa DNA/binding subunit). Slight decreases of interactions were observed with cytoplasmic NSF (vesicle-fusing ATPase) taking part in vesicle-mediated transport within the Golgi and from ER to Golgi, and endosomal SH3G1 (endophilin-A2), predicted to participate in endocytosis and recruitment of proteins to membranes.

## DISCUSSION

We report a large family with a distinct somewhat lupoid-like and SAVI-like phenotype carrying a novel G207E mutation in the substrate binding domain of STING. We show that the varying disease severity in this family is due to coding polymorphisms in *TMEM173* and *IFIH1*. The disease retained some typical clinical characteristics of SAVI such as cutaneous vasculitis, severe infections including necrotizing cellulitis, exacerbation of skin symptoms by cold and nasal septum defects (5, 11, 14). The condition was considerably milder and lacked lung manifestation, violaceus and ulcerative facial and acral skin lesions, elevated inflammatory markers, febrile attacks and autoinflammatory markers. (**Supplementary figure 1A-B**). The characteristic early-onset multisystemic inflammation was absent, and most patients did not develop life-threatening ulcers, did not require prolonged immunosuppressive medication and have had a normal lifespan. However, our patients presented with several previously unreported STING-associated lupoid-like features including alopecia, photosensitivity, and thyroid dysfunction. Most of them had a marked and widespread livedo reticularis from birth. Furthermore, their IFN score was less severely elevated.

Consistent with previous reports in SAVI patients (5, 25), all our patients had expanded population of naïve CD4+ and CD8+ T-cells but low numbers of mature B and T cells. The skewed maturation might be due to the chronic inflammation and immune cell exhaustion. STING has also been proposed to act as an inhibitor of immune cell proliferation (25), an effect linked to NFKB activation and spatially limited to Golgi (25). This finding is supported by our data, which shows hyperactivation of NFKB in the patient cells, and also an altered localization of the G207E mutant within the Golgi/ER compartment.

Since we and others (5, 11, 13-15) have seen considerable variation in disease severity among the *TMEM173* mutant carriers, we further analyzed the effect of polymorphisms in *TMEM173* (p.R232H) and *IFIH1* (p.A946T) on disease phenotype. The patients carrying the R232 variant had more severe disease. *In vitro,* this polymorphism strengthened the constitutive activation of the mutant STING, leading to the overexpression of STING downstream targets IFN, IL1B, and IL18. The observation is consistent with previously reported effect of R232 on amplifying IFNB pathway activation *in vitro* (6). We also noted an additive clinical effect for the G207E mutant and the two polymorphisms, with the combination R232+G207E+IFIH1 T/T leading to a severe earlyonset phenotype that resembles the SAVI-features (Pat V.2). *IFIH1* risk allele has been shown to promote disease through a combinatorial effect with other risk alleles (24). Although supported by experimental data, the additive effects will need validation in larger patient sets.

Understanding of the crosstalk between STING-pathway and the NLRP3 inflammasome controlling IL1B and IL18 secretion is rapidly expanding, and recently STING was shown to have a role in inflammasome activation (37, 38). We observed aberrant alternative NLRP3 inflammasome activation in response to bacterial TLR stimuli LPS and Pam3Cys in the patient PBMCs. The constitutively active STING-signaling in our patients’ PBMCs could explain the elevated LPS-induced IL1B secretion and drive the observed aberrant inflammasome activation in response to Pam3Cys as LPS-TLR-signaling and the following inflammasome activation utilizes an adaptor protein TRIF, that also promotes STING signaling (40, 53). In agreement with our data LPS was reported to aberrantly induce IL1B secretion in *TRIF*^*-/*^monocytic cells artificially expressing a SAVI-associated V155M STING mutant (37). Activation of NLRP3 inflammasome has a key role in the pathogenesis of other autoinflammatory diseases, such as cryopyrinopathies (54). Our data suggest that the STING GOF patients’ immune cells have a strong propensity for spontaneous inflammasome activation, provided that suitable inflammasome priming signals are present, and further imply that dysregulation of both interferons and IL1B may contribute to the disease phenotypes of interferonopathies.

To our knowledge, our study is the first to define the molecular interactions of STING. The majority of STING partners were Golgi, ER or endosomal proteins, consistent with reports of subcellular localization of STING (8, 33). The monogenic GOF mutants N154S and G207E led to an overall reduction of interactions. The most distinct differences between the classical N154S SAVI and the novel G207E mutation are the lost interactions with COPI and COPII components. These two protein complexes are essential for protein transport between ER and Golgi (55), and loss of COPI also causes monogenic autoimmune disease (52). The increased G207E interactions included Sequestosome-1 (SQSTM), which is an autophagy receptor involved in endosome organization (56). The combined loss of COPI and gained SQSTM interactions together with aberrant subcellular localization, verified with MS microscopy, suggest impairment in mutant STING trafficking. Since STING has been shown to predominantly signal from endosomes, the impaired trafficking likely leads to prolonged signaling and activation of downstream pathways (55, 57). Interaction with AP-1 complex subunit beta-1 (AP1B1), a protein involved in cargo sorting at trans-Golgi network, was lost or diminished in both H+207E and R+207E, further supporting the model of defective transport.

The interactomes of G207E in polymorphic backgrounds (H232 and R232) were highly similar. The R+G207E variant, which is associated with severe clinical disease, further diminished the G207E interactions with the COPI proteins-likely exaggerating the trafficking defect, and further prolonging STING signaling from the endosomes. R+207E variant also associated more with ubiquitin carboxyl-terminal hydrolase CYLD, which regulates inflammation and immune responses via NFKB-inducing pathways (58-60).

To conclude, we describe in a large multigenerational family a novel STING mutation G207E as a cause for a combination of SAVI-associated vasculopathy and selected lupoid-like features. The clinical heterogeneity is explained by *TMEM173* polymorphism p.R232H, and *IFIH1* risk variant p.A946T, with the two common variants showing cumulative effect on disease severity. We also describe the interactomes of wt and mutant STING, demonstrating a localization defect in the latter, and report elevated NRLP3 inflammasome activation and IL1B secretion in patient PBMCs, potentially contributing to the disease phenotype. The results broaden the clinical spectrum of STING mutation phenotypes, and increase our understanding of the molecular mechanisms of nucleic acid-driven inflammatory diseases.

## METHODS

### Patient information

From birth, the index case **<u>IV.1</u>** (**Figure 1**) has had erythema on the cheeks and severe livedo reticularis on his body and extremities, most pronounced on his sides, thighs, upper arms, hips and buttocks. Later, remitting idiopathic trombocytopenia and unspecified nail changes were noted at the age of 7y, as well as alopecia areata. Later, alopecia universalis with sparing of genital, eyebrow and some scalp hair has developed. Repeated bone marrow aspirates and karyotype have been normal. He has nasal septal perforation and slightly dysmorphic features with hypoplastic alae nasi and deep philtrum. He is sensitive to UV radiation and burns extremely easily. He was admitted to the dermatology ward at age 29y due to painful necrotic ulcerations resembling vasculitis on the shin and ankle that had appeared after a sunburn. Skin biopsy confirmed cutaneous vasculitis but extended laboratory tests were normal, except for an elevated Factor VIII (192-211 %), considered an independent risk factor for venous thrombosis **(Figure 1E).** His vasculitis was first treated with prednisolone (from 0.8mg/kg and tapered down) combined with azathioprine (150 mg, 1.5 mg/kg) or methotrexate (20 mg/week, 0.2 mg/kg), with poor response. Vasculitis then responded well to cyclosporine (maximum dose 250 mg/day, 2.5 mg/kg). A subsequent sunburn however induced re-flaring of previous ulcers. These subsided with topical corticosteroids. Additionally, he has had *Streptococcus pyogenes* periobital cellulitis and a single abscess on his inner thigh, but otherwise displays no susceptibility to infections.

IV.1 has three children, of whom daughter **<u>V.2</u>** shares a very similar phenotype. Since birth, she has had cheek erythema and severe livedo reticularis of extremities and trunk, most severe on her thighs, upper arms and sides. She has had UV-induced urticaria. At age 6.8y, she developed alopecia areata, which progressed to universal alopecia with loss of all body hair except the eyebrows and eyelashes by age 7.8y. At age 12.5y she was diagnosed with autoimmune thyroiditis (TPOAb 370 IU/mL) and hypothyroidism. She has a nasal septal defect and similar dysmorphic features as her father. At age 3y, she had *Streptococcus pyogenes* sepsis and necrotizing cellulitis on her upper arm and abdomen. At age 10y, she developed an eschar on her abdomen, which developed rapidly into a large necrotic abscess with *Streptococcus pyogenes* **(Figure 1C).** Surgical revision of the necrotic and infected tissue produced a large wound on the abdomen **(Figure 1D).** Skin biopsy from the edge of the necrosis was indicative of small vessel vasculitis and underlying fasciitis. She was successfully treated with i.v. antibiotics, prednisolone (gradually tapered down from 0.5 mg/kg) and cyclosporine 100 mg/d (2.1 mg/kg). Treatment with negative pressure wound therapy successfully assisted wound closure.

From birth, the mother of IV.1 (**<u>III.4)</u>**, has had mild livedo reticularis on the buttocks, thighs and upper arms. At age 7y she developed alopecia areata and was diagnosed with hypothyroidism. She burns easily in the sun and has constant facial erythema.

**<u>III.7</u>** developed livedo reticularis first on the buttocks at age 8y, from where it spread to thighs and upper parts of shins, upper arms, shoulders and stomach, and most recently on her breasts. She also has facial erythema. The livedoid discoloration continuously spreads and increases. She burns easily and had UV intolerance as a child. She had early menopause at age 42y and since then has suffered from severe periodontitis.

**<u>IV.6</u>** has had mild livedo reticularis form early childhood on her upper arms and knees. She burns easily in the sun. At age 10y she developed total alopecia and, has since lacked all body hair while her scalp hair regrew within a few months. At age 12y, total alopecia reoccurred. She has occasionally also had partial loss of eyelashes. At age 13y, she was diagnosed with autoimmune thyroiditis and hyperthyroidism. Anti-thyroid peroxidase-antibody (TPOAb) titers reached 1855 IU/mL at age 21y. She was then also diagnosed with pituitary thyroid resistance and received radioiodine treatment. Thereafter, she has received thyroid hormone substitution. She has constantly elevated calcitriol and calcium levels with normal parathyroid hormone levels, not indicative of vitamin D resistance. However, after initiating calcium and vitamin Dreplacement therapy, she has had minor regrowth of scalp hair. She has recurrent periodontitis and tooth enamel defects. Despite exercise-induced shortness of breath, lung function tests (spirometry and diffusion capacity) were normal.

From adulthood, **<u>III.9</u>** has had severe livedo reticularis, which slowly progressed on her upper arms, posterior armpits, sides, lateral back region and buttocks extending to posterior thighs. At age 4y, she experienced total alopecia with subsequent hair regrowth. During her childhood and adolescence, she had recurrent, intermittent spells of alopecia areata. At age 30y, total alopecia with sparsening of eyelashes reoccurred and a vitiligo spot appeared on her cheek. At age 38y, she was diagnosed with autoimmune thyroiditis with TPOAb levels >1000 IU/mL. She has suffered from infertility that is unresponsive to treatment. She has a nasal septal defect, but is not sensitive to UV light. She has had recurrent sinusitis and sleep apnea, while lung function tests were normal.

A mild clinical immunodeficiency in the family was suggested by the severe and persistent periodontitis, recurrent upper respiratory tract and skin infections (**Table 1**). Lupoid and autoimmune features included photosensitivity and erosion of the nasal septum (**Table 1**). However, no antinuclear, anti-DNA or other lupus autoantibodies were detected (**Supplementary table 1**). Most (5/6) patients had elevated serum IgE levels (**Supplementary table 1**).

### DNA extraction

Genomic DNA was extracted from EDTA blood samples or salivary samples using Qiagen FlexiGene DNA kit (Qiagen, Valencia, CA, USA) or OraGene DNA Self-Collection Kit (OGR-250, DNA Genotek, Holliston, MA, USA).

### Genotyping and linkage analysis

The DNA samples of 10 family members were run on the IlluminaHumanOmniExpressExome-8v1-2 array at the Science for Life Laboratory SNP&SEQ platform in Uppsala, Sweden, according to standard protocols (Illumina, San Diego, CA, USA) and the results were analyzed using the software GenomeStudio 2011.1 from Illumina Inc. Two controls were run in parallel. Genotyping was based on cluster files generated from the signal intensities from more than 800 DNA samples processed in parallel to this project.

Samples and markers with low call rates were removed (<95% vs <99%, respectively), and the remaining markers were checked for inheritance errors using PedCheck (61). LD-based pruning of the genotype data was performed using the pairwise LD threshold function in PLINK (v1.07 (62)), (--indep-pairwise 50 5 0.2).

The Rutgers genetic map interpolator Rutgers Map v2 (63) was used to attain genetic map information for the markers in the linkage analysis. Merlin (v1.1.2 (64)) was used to perform parametric and non-parametric linkage analysis, using the LD-pruned genotype dataset.

### Whole genome sequencing

Next generation sequencing libraries were built according to established laboratory protocols at the Science for Life Laboratory Stockholm and the Institute for Molecular Medicine Finland (FIMM). The exome libraries were processed according to the Agilent SureSelect Target Enrichment System (Agilent Technologies, Santa Clara, CA, USA) for Illumina Paired-End Sequencing Libraries (Illumina, San Diego, CA, USA) using the SureSelect Human All Exon V5 capture library (Agilent Technologies, Santa Clara, CA, USA). Libraries were sequenced with 101 bp read length (HiSeq1500 sequencing platform, Illumina, San Diego, CA, USA).

The read mapping, variant calling and genome annotation were performed as described previously (65). We focused the search on the haplotype that segregated with the phenotype, prioritizing rare, heterozygous coding variants that negatively affect conserved residues and shared by both whole genome sequenced individuals. Only one such variant was identified, localized within the *TMEM173* gene. The G207E variant was absent from all major public (dbSNP, Exome Aggregation Consortium) and in-house genome databases.

### Verification of candidate mutations

The candidate mutations were verified by capillary sequencing from blood DNA samples using BigDye chemistry (Terminator v3.1, Life Technologies, Carlsbad, CA, USA) and the TMEM173 p.(Gly207Glu) forward 3’CCAATGACCTGGGTCTCACT-5’, and reverse 3’AAGCTCATAGATGCTGTT-GCTG-5’ primers.

### Targeted TMEM173 PCR amplicon sequencing

Sample preparation was performed according to an in-house targeted PCR amplification protocol. All oligonucleotides were synthesized by SigmaAldrich (St. Louis, MO, USA). The protocol includes two rounds of PCR amplification. The first round of amplification was performed in a volume of 20 µl containing 10 ng of sample DNA, 10 µl of 2x Phusion High-Fidelity PCR Master Mix (Thermo Scientific Inc., Waltham, MA, USA), 0,25 µM of each locus specific primer carrying Illumina-compatible adapter sequences (**Supplementary table 2**), and the reaction mix was brought to a final volume with laboratory-grade water. Cycling was performed according to Phusion High-Fidelity PCR Master Mix cycling instructions (annealing temperature 59 °C). Samples were purified with Performa® V3 96-Well Short Plate (EdgeBio, Gaithersburg, MD, USA) and QuickStep™2 SOPE™ Resin (EdgeBio, Gaithersburg, MD, USA) according to the manufacturer’s instructions. Subsequent to the purification, a second round of PCR amplification was performed in a volume of 20 µl, containing diluted PCR product from the first round of PCR. 10 µl of 2x Phusion High-Fidelity PCR Master Mix (Thermo Scientific Inc., Waltham, MA, USA), 0,375 µM of both index primers carrying Illumina grafting P5/P7 sequence and the reaction mix was brought to a final volume with laboratory grade water. Samples were cycled as follows: initial denaturation at 98 °C for 2 minutes; 10 cycles at 98 °C for 20 seconds, at 65 °C for 30 seconds and at 72 °C for 30 seconds; final extension at 72 °C for 5 minutes, followed by a hold at 10 °C.

Following PCR amplification, samples were pooled in equal volumes. The sample pool was purified twice with Agencourt® AMPure® XP (Beckman Coulter, Brea, CA, USA) according to the manufacturer’s instructions using equal volumes of the Agencourt® AMPure® XP and the sample pool. The purified sample pool was analyzed on Agilent 2100 Bioanalyzer using Agilent High Sensitivity DNA Kit (Agilent Technologies Inc., Santa Clara, CA, USA) to quantify amplification performance and yield.

Sequencing of PCR amplicons was performed using Illumina MiSeq instrument with MiSeq Control Software v2.5 (Illumina, Inc., San Diego, CA, USA). Samples were sequenced as 251 bp paired-end reads and two 8 bp index reads. The read mapping, variant calling and genome annotation were performed as described previously (65).

### Peripheral blood immunophenotyping

Fresh EDTA-blood samples or PBMNCs were used for B and T lymphocyte immunophenotyping. Four or 6-color flow cytometry panel with mAbs against the surface antigens IgM, IgD, CD3, CD4, CD8, CD16 /56, CD19, CD21, CD27, CD33, CD34, CD38, CD45, CD56, CD57, CD133, HLA-DR, CD62L, CD45RA and CD45RO (BD Biosciences, Franklin Lakes, NJ, USA) was used (65). The memory status of T cells was studied with the antibody panel including anti-CD45, -CD3, -CD4, -CD45RA, and -CCR7 (R&D Systems, Minneapolis, MN, USA) (65, 66).

Evaluation of T cell responses is described in detail elsewhere (66). For the assessment of T cell activation, fresh mononuclear cells were stimulated for 6 hours with anti-CD3, anti-CD28, and anti-CD49d (BD Biosciences, Franklin Lakes, NJ, USA). The cells were analyzed using a 4or 6-color flow cytometry panel with mAbs against the antigens CD45, CD3, CD4, CD8, CD16, CD56, CD45, CD45RA, TCR-γ, CCR7, IFN-γ, and tumor necrosis factor (TNF).

### Site-directed mutagenesis predictions

The 3D structures of STING (4LOI and 4KSY; (32, 67)) were extracted from Protein Data Bank (PDB; www.rcsb.org, (68)) and the effects on the protein stability of the individual mutants analyzed with SDM (26).

### Construct generation

The *TMEM173* cDNA was obtained as a gateway compatible entry-clone from human Orfeome collection (Horizon Discovery, UK). This entry-clone has the p.His232, which is the minor allele in population. All *TMEM173* variant constructs were created using Q5 Site-directed Mutagenesis Kit (New England Biolabs, Ipswich, MA, USA). The mutagenesis primers are listed in **Supplementary table 2**. All the generated variant entry-constructs were confirmed with direct sequencing prior to introduction to C-MAC-Tag destination vector (45).

### Luciferase assay

Human embryonic kidney (HEK293) cells were cultured at 37°C in 5% CO_2_ in DMEM (SigmaAldrich, Espoo, Finland) supplemented with 10% FBS (Sigma), 50 mg/ml penicillin, and 50 mg/ml streptomycin. 20 000 cells were seeded onto Costar 3610 white, clear bottom 96-well plates, and the following day co-transfected with 50 ng of TMEM173 constructs or empty C-MAC-tag-vector (45), 40 ng of IFNB promoter-driven firefly luciferase reporter plasmid (IFN-β-pGL3; a kind gift from Yanick J. Crow), and 1.4 ng Renilla luciferase reporter plasmid (pRL-SV40) by using FuGENE 6 (Promega, Madison, WI, USA). Transfected cells were stimulated 24 hours later with cyclic dinucleotides by transfecting 4 µg/ml 2’3’-cyclic guanosine monophosphate (cGAMP), or 3’3’cGAMP or c-di-GMP (all Invivogen, San Diego, CA, USA) with Lipofectamine 2000 (Thermo Fischer Scientific, Vantaa, Finland). After 24 hours incubation, the cells were lysed with 1X passive lysis buffer (Promega). Luciferase assays were performed with the Dual-Glo Luciferase Assay System (Promega) according to manufacturer’s instructions. IFN-β-pGL3 plasmid was utilized to measure IFN- promoter activity (firefly luciferase), and Renilla luciferase plasmid was used for transfection efficiency normalization. For final results, counts of untransfected cells were subtracted from both firefly and Renilla luciferase, and firefly luciferase activity further normalized against Renilla luciferase activity.

### Nanostring analysis of patient PBMCs

Blood samples were collected from patients (V.2, IV.1, III.4, III.7) and matched controls in Vacutainer® CPT™cell preparation tubes with sodium heparin^N^ (BD, New Jersey, USA) and PBMCs were isolated as instructed by manufacturer. Approximately 10 million PBMCs for each patient were pelleted, snap frozen and stored at -80 °C until RNA extraction. RNA was extracted with the RNeasy Mini Kit (Qiagen, Hilden, Germany) according to manufacturer’s protocol and concentration measured with a Qbit fluorometer (Thermo Fisher Scientific, WA, USA). For each sample, 100 ng RNA in 5 µl volume was pipetted into the NanoString nCounter gene expression platform (NanoString Technologies, Seattle, USA). Our custom gene set containing 50 genes includes IFN-regulated, inflammasome-related, as well as JAK/STAT and NFKB signaling pathway genes. For overnight hybridization at 65 °C, RNAs were mixed with 5’ reporter probes (tagged with fluorescent barcodes of targeted genes) and 3’ biotinylated capture probes. Reactions were purified and immobilized on the sample cartidge surface by utilizing the Prep Station (NanoString Technologies), and the cartridge was scanned in triplicate using an nCounter Digital Analyzer (NanoString Technologies). Analysis of gene expression data was performed with the nSolver™ 4.0 analysis software (NanoString Technologies). As recommended by the manufacturer, we performed two-step normalization. First, background thresholding using negative control counts setting threshold to mean +2 standard deviations above the mean was used. Then, second, to remove technical variability we utilized positive control and CodeSet content (housekeeping gene) two-step normalization. Both normalizations were performed using default settings unless otherwise stated. The normalized values were utilized to calculate interferon scores. Interferon scores were calculated similarly to (36). Briefly, the fold change for each individual gene between patient and matched control was calculated, and the median fold change of all was used to calculate the interferon score for each patient.

### Culture and stimulation of peripheral blood mononuclear cells (PBMCs)

PBMCs were isolated from fresh EDTA-anticoagulated blood of patients (IV.1, III.4, III.7) and sex- and age-matched control subjects by density gradient centrifugation in Ficoll-Paque PLUS (GE Healthcare, UK). After four washes in PBS, the cells were seeded in 24-well culture plates at 1.5 x 10^6^ cells/ ml in RPMI 1640 (Lonza, Basel, Switzerland) supplemented with 2 mM GlutaMAX (Gibco, Thermo Fischer Scientific, Finland), 100 U/ml penicillin -100 µg/ml streptomycin (Gibco), and 10 % fetal bovine serum (Gibco). The cells were allowed to rest for a minimum of 3 h and subsequently stimulated with 1 μg/ml lipopolysaccharides (LPS) from *E.coli* O111:B4 (Sigma), 1 μg/ml Pam3Cys-SKKKK (Pam_3_Cys) (EMC microcollections), and 5 mM ATP (stock solution neutralized; Sigma) for the indicated times.

### Measurement of cytokine secretion

TNFA and the mature, cleaved forms of IL1B and IL18 were detected from PBMC culture media supernatants using Human TNFA DuoSet ELISA, Human IL1B/IL1F2 DuoSet ELISA, and Human Total IL18 DuoSet ELISA (all from R&D Systems).

### Quantitative real-time PCR

PBMC RNA was isolated using the RNeasy Plus Mini Kit (Qiagen), followed by cDNA synthesis with iScript kit (Bio-Rad). Quantitative real-time PCR was run in duplicate reactions of 10 ng cDNA using LightCycler480 SYBR Green I master (Roche Diagnostics, Espoo, Finland) and a LightCycler96 instrument (Roche). See **Supplementary table 2** for primer sequences. Relative gene expression was calculated using the 2^(-ΔΔCt)^ method. The data was normalized against geometric mean of expression of two housekeeping genes (ribosomal protein lateral stalk subunit P0 and beta-2-microglobulin).

### Cell culture

To generate stable, isogenic, inducible cell lines Flp-In™ T-REx™ HEK293 cells (Invitrogen, Life Technologies) were transfected with the generated STING-MAC-tagged constructs using FuGENE6 (Promega) as described earlier (69). Cells were cultured as instructed by manufacturer.

Five 14 cm plates for each construct and biological replicate were induced either with 1 µg/ ml tetracycline and 50 µM biotin 24 hours prior to harvesting. Harvesting, cell lysis, affinity-purification, mass spectrometry were performed as described (45, 69).

### Mass spectrometry data analysis and filtering

The mass spectrometric data was searched with Proteome Discover 1.4 (Thermo Fischer Scientific) using the SEQUEST search engine against the UniProtKB/SwissProt human proteome (http://www.uniprot.org/, version 2015-09). Search parameters were set as in (45). All data were filtered to highconfidence peptides according to Proteome Discoverer FDR 1%. The lists of identified proteins were conventionally filtered to remove proteins that were recognized with less than two peptides and two PSMs. The high-confidence interactors were identified using SAINT and CRAPome as in (45). Each sample’s abundancy was normalized to its bait abundancy. These bait-normalized values were used for data comparison and visualization.

### Mass spectrometry microscopy (MS microscopy)

The bait-normalized values from BioID data obtained from previous step were used as input for MS microscopy analysis (http://www.biocenter.helsinki.fi/bi/protein/msmic.helsinki.fi/bi/protein/msmic) (45). MS microscopy calculates a localization score that numerically describes the bait protein’s dynamic localization in cell.

### Statistics

Statistical analysis using 2-tailed Student’s t-test was performed using Microsoft Excel. A p-value less than 0.05 was considered significant.

### Study approval

The study was conducted in accordance to the principles of the Helsinki Declaration and was approved by the Helsinki University Central Hospital Ethics Committee (91/13/03/00/2011). Written informed consent was obtained from the patients and healthy controls.

## Author contributions

SK, EH, EE, KR designed and performed experiments, analyzed data, wrote the manuscript and prepared the figures. HH, MI, MP, EM, KH, and SL performed experiments. SKi, SM, KE, JS, and JK supervised experiments and data analysis. MS, AR, KH-J, and MV designed experiments and wrote the manuscript.

### Acknowledgements

We sincerely thank the patients for their co-operation and patience. Eira Leinonen, Auli Saarinen and Alli Tallqvist are acknowledged for their technical assistance. Sini Miettinen is acknowledged for her excellent assistance with MS operation. Xiaonan Liu and Vera Varis are thanked for excellent technical assistance in the lab. PhD Shabih Shakeel and PhD James Geraets are thanked for their advice and critical comments on the SDM predictions. PhD Tiina Öhman is thanked for critical review of the manuscript. The Institute for Molecular Medicine Finland (FIMM) Technology Centre, Helsinki and Science for Life Laboratory, Stockholm are acknowledged for providing research infrastructure services. The authors acknowledge support from Science for Life Laboratory, the Knut and Alice Wallenberg Foundation, the National Genomics Infrastructure funded by the Swedish Research Council, and Uppsala Multidisciplinary Center for Advanced Computational Science for assistance with massively parallel sequencing (alternatively genotyping) and access to the UPPMAX computational infrastructure. Genotyping was performed by the SNP&SEQ Technology Platform in Uppsala. The platform is part of Science for Life Laboratory at Uppsala University and supported as a national infrastructure by the Swedish Research Council. The Academy of Finland, Helsinki University Hospital Research Funds, Finnish Medical Foundation, Foundation for Paediatric Research, Sigrid Juselius Foundation, Finska Läkaresällskapet, Finnish Cancer Institute, Karolinska Institutet Research Foundation, Swedish Research Council and Strategic Research Programme in Diabetes, Karolinska Institutet supported this work.

---

**Supplementary figure 1:**
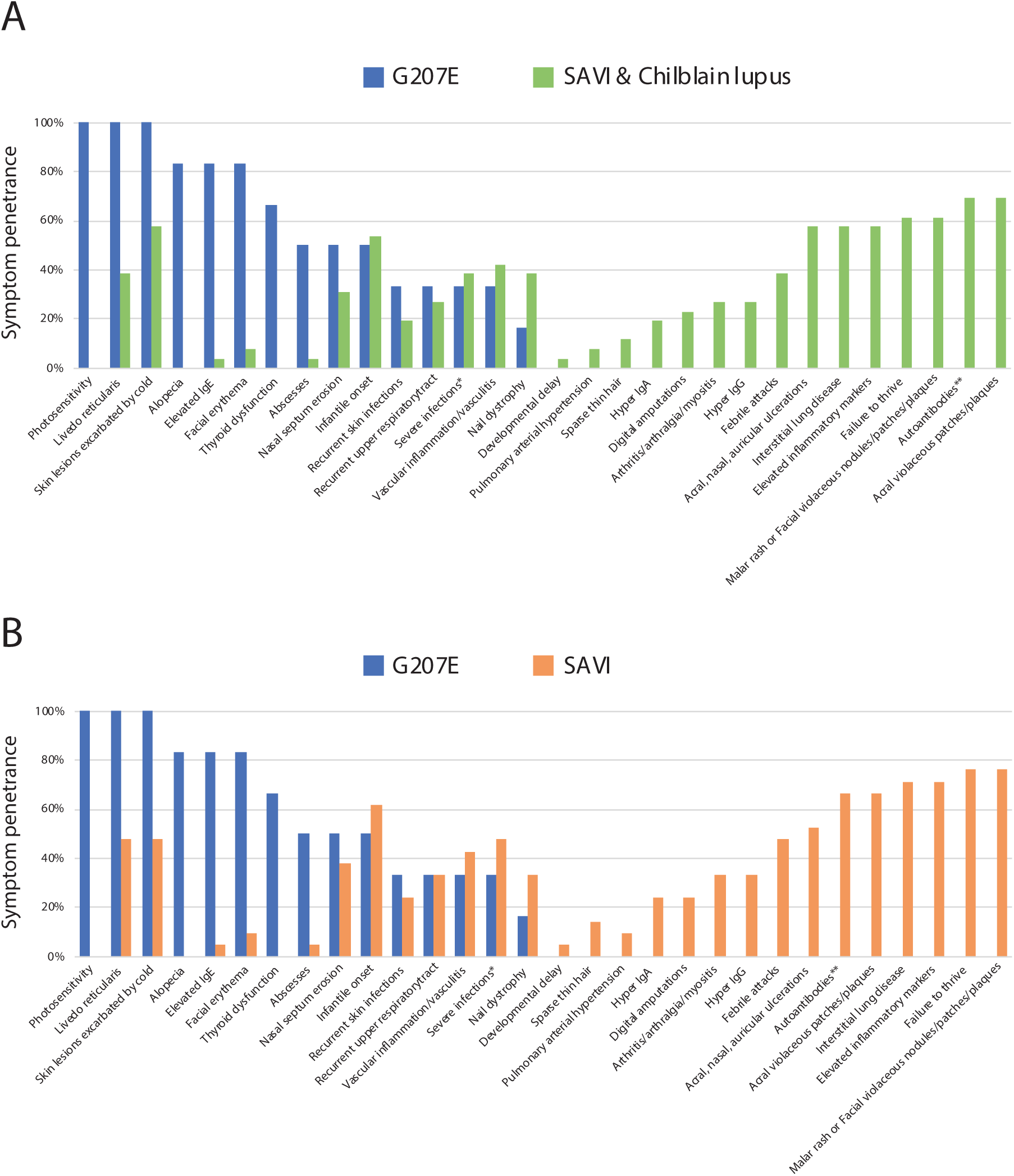
A) Comparison of clinical phenotypes in G207E mutation carriers (n=6) and in previously reported SAVI (n=21) (5, 11, 12, 14, 15, 70-73) and chilblain lupus patients (n=5) (13). * Pneumonia, cellulitis, necrotizing fasciitis, septicemia; ** Antinuclear 54%, antiphospholipid 19%, ANCA 15%, cardiolipin 4% B) Comparison of G207E carriers (n=6) and SAVI patients (n=21). * Pneumonia, cellulitis, necrotizing fasciitis, septicemia; ** Antinuclear 48%, antiphospholipid 24%, ANCA 19%, cardiolipin 5%

**Supplementary figure 2:**
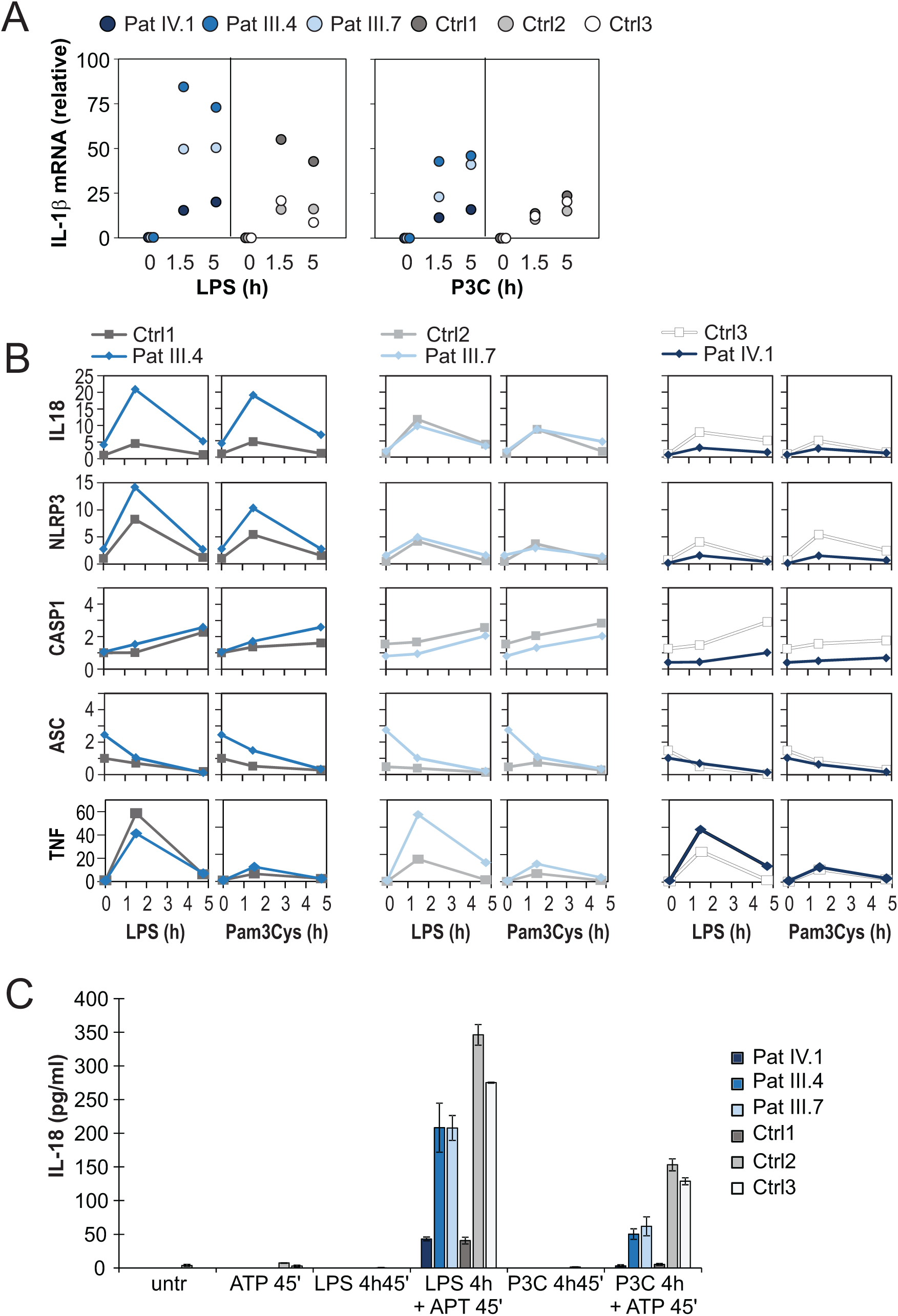
A-B) Inflammasome-related mRNA expression levels, and C) amount of mature IL18 in PBMC culture supernatant after activation of canonical NLRP3 pathway by short TLR priming followed by 45 min ATP stimulation.

**SUPPLEMENTARY TABLE 1:** Blood counts, autoantibodies and functional assay findings in STING G207E carriers.

| Patient | Control Range | IV.1 | V.2 | III.4 | III.7 | III.9 | IV.6 |
| --- | --- | --- | --- | --- | --- | --- | --- |
| **Leukocytes** (10<sup>6</sup>/l) | 3400-8200 | 4400 | 5100 | 4500 | 5900 | 5600 | 4200 |
| **Lymphocytes** (10<sup>6</sup>/l) | 1300-3600 | 2240 | 1880 | 1600 | 2180 | 1860 | 1270 |
| **Monocytes** (10<sup>6</sup>/l) | 200-800 | 190 | 110 | 540 | 500 | 220 | 100 |
| **Neutrophils** (10<sup>6</sup>/l) | 1500-6700 | 1820 | 2840 | 2190 | 3040 | 3200 | 2580 |
| **Basophils** (10<sup>6</sup>/l) | 0-100 | 20 | 10 | 10 | 10 | 10 | 0 |
| **Eosinophils** (10<sup>6</sup>/l) | 30-440 | 120 | 270 | 180 | 120 | 300 | 100 |
| **B cells (CD19+)** (10<sup>6</sup>/l) | 100-500 | 70 | 340 | 180 | 180 | 290 | 50 |
| **CD3+CD4+** (10<sup>6</sup>/l) | 300-1400 | 802 | 929 | 869 | 1065 | 1030 | 712 |
| **CD3+CD8+** (10<sup>6</sup>/l) | 200-1200 | 290 | 310 | 250 | 600 | 430 | 210 |
| **NK cells (CD3-CD16<sup>+</sup>56<sup>+</sup>)** (10<sup>6</sup>/l) | 90-600 | 50 | 180 | 180 | 280 | 120 | 200 |
| **Platelets** (10<sup>6</sup>/l) | 150 000-360 000 | 107 000 | 254 000 | 229 000 | 279 000 | 274 000 | 173 000 |
| **ANA-ab** (titer) | <320 | <80 | <80 | <80 | <80 | <80 | <80 |
| **ENA-ab** | Negat | Negat | Negat | Negat | Negat | Negat | Negat |
| **DNA-ab** (IU/ml) | <10 | <10 | <10 | NA | <10 | <10 | <10 |
| **ANCA** | Negat | Negat | Negat | Negat | NA | Negat | NA |
| **LupusAK** | Negat | Negat | Negat | NA | Negat | Negat | NA |
| **PLAb** | Negat | Negat | NA | NA | Negat | NA | NA |
| **KardAbG** (U/ml) | <15 | <4 | <4 | NA | <4 | 4 | NA |
| **IgE** (kU/l) | 0-110 | 290 | 4418 | 738 | 889 | 46 | 889 |
| **Complement** | Classical, alternative, mannan-binding lectin pathway hemolytic activities | Normal | Normal | Normal | Normal | Normal | Normal |
| **Lymphocyte proliferative responses to mitogens** | Phytohemagglutinin, concanavalin A, pokeweed mitogen | Normal | Normal | NA | NA | Normal | NA |

Negat= negative, NA = not available.

**SUPPLEMENTARY TABLE 2:**
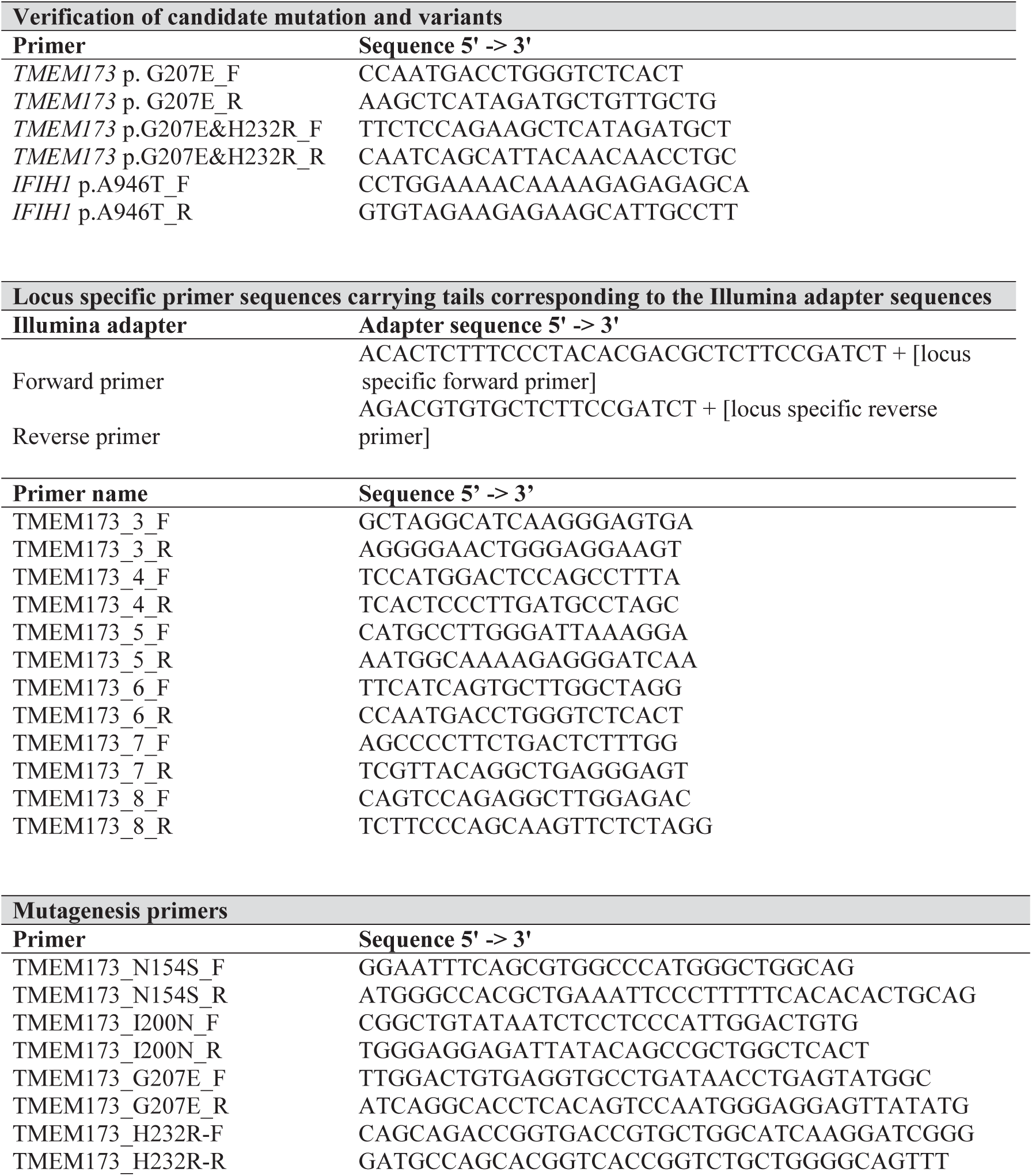
Primer list. (F, forward primer; R, reverse primer)

## Abbreviations

2’3’-cGAMP: cyclic GMP-AMP, mammalian cGAMP (tlrl-nacga23)
3’3’-cGAMP: cyclic GMP-AMP, bacterial cGAMP (tlrl-nacga)
c-di-GMP: cyclic di-guanylate monophosphate (tlrl-nacdg)
CDN: cyclic-di-nucleotide
FCL: familial chilblain lupus
GOF: gain-of-function
IFIH1: interferon induced with helicase C domain 1
SAVI: STING-associated vasculopathy with onset in infancy
SLE: systemic lupus erythematosus
SMS: Singleton-Merten syndrome
STING: Stimulator of interferon genes protein
TMEM173: transmembrane protein 173

## Notes

**CONFLICT OF INTEREST**: M.S. has received honoraria from CSL Behring. S.M. has received honoraria from BMS and Novartis. J.S. has received honoraria from Roche. K.H-J. has received honoraria from Octapharma and Abbvie. A.R. is an Advisory Board Member of ImmunoQure Gmbh. Other authors declare no conflict of interest.

